# Modulating adipose-derived stromal cells’ secretomes by culture conditions: effects on angiogenesis, collagen deposition, and immunomodulation

**DOI:** 10.1101/2024.11.01.621486

**Authors:** Erika Pinheiro-Machado, Claudia C. Koster, Alexandra M. Smink

## Abstract

The secretome of adipose-derived stromal cells (ASC) presents a promising avenue for cell-free therapies due to their rich mixture of bioactive molecules. Different culture conditions can modulate the composition of this mixture, but how this affects the functional properties of the secretome remains to be investigated. This study investigated the *in vitro* effects of normoxia, cytokines, high glucose, hypoxia, and hypoxia + high glucose-derived ASC secretomes on angiogenesis (tube formation assay), collagen deposition (Picrosirius-Red staining), and immunomodulation (One-way Mixed Lymphocyte Reaction in combination with an antibody-mediated cell-dependent cytotoxicity assay). The data showed that normoxia and hypoxia-derived secretomes consistently exhibited potent proangiogenic effects in both human and rat models. These secretomes also demonstrated positive influences on collagen deposition and immunomodulation. Interestingly, the human ASC hypoxia + high glucose-derived secretome emerged as a stimulator of collagen deposition and modulator of the immune system. Conversely, cytokines and high glucose-derived secretomes have shown less strong effects in almost all functional parameters. In conclusion, our findings indicate that modulating culturing conditions results in secretomes with different functional properties and emphasizes the multifaceted role of ASC secretomes in regenerative processes.

## Introduction

Adipose-derived stromal cells (ASC) are adult stem cells found in various adipose tissue depots (1, 2). Due to these cells’ ability to differentiate into multiple lineages and self-renew (3, 4), extensive research into their regenerative potential has been performed (3, 5). Investigation of ASC cell-based approaches showed that these cells could, among others, modulate the immune system (5–7), stimulate angiogenic processes (8–10), influence extracellular matrix (ECM) remodeling (11–13), and support cell survival (3, 14). These inherent features make ASC a promising cell type for regenerative medicine and tissue engineering applications.

Despite great promise, ASC cell-based approaches have challenges, limitations, and potential risks (3, 15, 16). Protocols for ASC isolation, expansion, and characterization are not yet fully standardized, and the research that has been done may encounter reproducibility issues as different ASC subpopulations may have different therapeutic outcomes (16–19). Once isolated, expanded, and characterized, ASC used in transplantation strategies display poor cell survival, undergoing cell death in or on the way to the target tissue due to inadequate vascularization and unfavorable microenvironments (20–22). Moreover, limitations related to the choice of the cells’ donors’ characteristics (such as age and metabolic profile), as well as the ASC administration route for the transplantation, are still to be resolved (3, 23, 24).

Where at first, the ASC regenerative capacity was thought to be dependent on the presence of these cells at the injury site, ASC secretome-based therapies have shown to harness these cells’ regenerative abilities, offering an alternative approach (25–27). The ASC secretome is a mixture of secreted factors that contains a variety of growth factors, cytokines, ECM proteins, exosomes, and more (16, 27, 28). This mixture can be modulated by applying different ASC cell culture conditions (29–31), thereby fine-tuning its regenerative value. Moreover, the use of the secretome offers important advantages, such as the reduced risk of cell-related adverse events, but also the simplified and multi-varied possibility for delivery methods, as well as the potential for different secretome formulations, and large-scale, standardized production (16, 27, 32, 33). Therefore, ASC secretome-based therapies hold great promise in regenerative medicine and are being explored as a safer and more scalable alternative to ASC transplantation (27, 34, 35).

We have previously shown that the composition of human and rat perirenal ASC (prASC) secretomes can be modulated upon exposure of prASC to different culturing conditions (36). *In-silico* analysis of these different secretome formulations revealed that these mixtures were enriched in various factors that participate in different pathways and processes crucial to health and disease (36). More specifically, we demonstrated that prASC secretomes resulting from the prASC exposure to normoxia, cytokines, high glucose, hypoxia, and hypoxia + high glucose are mostly enriched in pathways associated with blood vessel formation, ECM organization and the immune system (36) – all essential for a balanced and orchestrated regenerative process.

Here, we aimed to investigate the *in vitro* functional effects of these various prASC secretomes on the formation of tube-like structures, collagen deposition, and modulation of humoral alloimmunity as a readout for the potential involvement of these secretomes in the pathways highlighted in our previous *in-silico* results. prASC secretomes were generated under normoxia, cytokines, high glucose, hypoxia, and hypoxia + high glucose exposure, and they were used in a tube formation assay (TFA), Picrosirius-Red staining, and mixed lymphocyte reaction (MLR). These experiments provided insight into which specific culturing conditions and resulting secretome compositions can enhance or modulate these cellular processes. Ultimately, this research holds promise for advancing our understanding of ASC secretomes and their potential clinical applications in tissue engineering, angiogenesis-related therapies, and immune modulation strategies.

## Materials and methods

### Media preparation

This research used multiple media types: 1: standard medium without serum (STD-M (-)), which is used for adipose tissue digestion and for both human and rat prASC culturing upon normoxia, exposure to cytokines, and hypoxia, and it contains Dulbecco’s Modified Eagles Medium 4.5 g/L D-glucose (DMEM; Lonza, Walkersville, MD, USA) supplemented with 50 U/mL penicillin, 50 mg/L streptomycin (Corning Cellgro, Manassas, VA, USA), and 2 mM/L L-glutamine (Lonza); 2: standard medium with serum (STD-M (+)) which is used for both human and rat prASC isolation and expansion; culture of human fibroblasts, and isolating rat fibroblasts; control for the antibody-mediated cytoxicity (CDC) assay, and it contains DMEM 4.5 g/L D-glucose supplemented with 50 U/mL penicillin, 50 mg/L streptomycin, 2 mM L-glutamine, and 10% Fetal Bovine Serum (FBS; Thermo Fisher Scientific, Bleiswijk, The Netherlands); 3: high glucose medium (HG-M (-)), which is used for both human and rat prASC secretome harvesting upon culturing in high glucose and in hypoxia + high glucose, and it contains DMEM 35mM D-glucose, supplemented with 50 U/mL penicillin, 50 mg/L streptomycin, and 2 mM L-glutamine; 4: cytokine medium (Cyto-M (-)),which is used for human prASC secretome harvesting upon cytokine exposure and contains DMEM 4.5 g/L D-glucose, supplemented with 50 U/mL penicillin, 50 mg/L streptomycin, 2 mM L-glutamine, 20 ng/mL IFN-γ (ImmunoTools, Friesoythe, Germany), 21.5 ng/mL TNF-α (ImmunoTools), and 10 ng/mL IL-1β (ImmunoTools). Cyto-M (-) used for culturing rat prASC contained the same cytokines’ concentrations but used rat cytokines (ImmunoTools); 5: DMEM/F12, which was used for the human fibroblasts (FLF92) culturing and expansion and contains DMEM/F12 4.5 g/L D-glucose, 100 IU/mL penicillin, 20% FBS, 0.5 ug/mL gentamicin amphotericin B (Sigma-Aldrich), 100 µg/L streptomycin, and 2 mM L-glutamine; 6: endothelial cell growth basal medium (EBM-2 (+)) which was used to culture and expand HUVEC and it contains endothelial cell growth basal medium (EBM-2; Lonza, Allendale, United States) supplemented with EGM-2 MV SingleQuot Kit Supplements and Growth Factors (Lonza); 7: complete RPMI (RPMI (+)), which was used for the rat splenocytes isolation.

### Isolating and culturing prASC

#### Collection of adipose tissue

Human perirenal adipose tissue was obtained from living kidney donors at the University Medical Center Groningen (UMCG), as previously described (36). The samples used in this study were acquired through anonymous donations, with the informed consent of the involved individuals. The ethical board of the UMCG approved this process, adhering to the established guidelines for handling waste materials. The study included male (43%) and female (57%) human donors with an average age of 57.6 ± 11.5 years. The samples were stored at a temperature of 4°C and processed within 48 hours (h). In addition, perirenal adipose tissue from male Sprague Dawley rats (Envigo, Horst, The Netherlands), aged 7-9 weeks and weighing between 250 and 280 grams, was obtained. The collection of adipose tissue from rats was conducted following the approval of the Dutch Central Committee on Animal Testing (CCD) and Animal Welfare Authority at the University of Groningen (AVD10500202115138). All animals were housed at the Central Animal Facility of the UMCG during the experiment.

#### Processing adipose tissue and culturing prASC

Human (h-prASC) and rat (r-prASC) perirenal adipose-derived stromal cells were isolated from the collected adipose donor tissue. The isolation procedure followed the same steps for both species, and the procedure has been previously described (36). Briefly, the collected tissue was washed, cut into small pieces, and digested using collagenase neutral protease blend 4 (0.5 mg/mL; Nordmark Biochemical, Uetersen, Germany) in STD-M (-) at 37 °C while shaking for 30 minutes (min). Centrifugation (700 x g, 7 min, room temperature (RT)) separated the cells from fibrous material, adipocytes, and lipids. The resulting pellet, containing prASC, was resuspended in STD-M (+) in a T25 flask for initial cell culture [passage 0] at 37 °C and 5% CO_2_. Following 24 hours, the medium was removed, and cells were washed with phosphate-buffered saline (PBS; Thermo Fisher Scientific). The growth medium (STD-M (+)) was refreshed every three days. Once the cells reached 80% confluence, they were passaged. Cells in passages 3 – 5 were used to collect the secretomes used for the functional assays. Cells from both human and rat donors (passage 3, n = 3) were characterized by the presence of mesenchymal stromal cells (MSC) surface markers (immunophenotyping), differentiation, and stemness potential. Characterization data can be found in Pinheiro-Machado, E. *et al.* (36).

### Secretome harvesting

To obtain the various secretomes, h-prASC and r-prASC (passage 4, n = 3) were seeded in 6-well plates at a density of 50,000 cells per well and cultured in STD-M (+). Once the cells reached 80% confluence, they were carefully washed with PBS and replenished with STD-M (-). After overnight serum starvation, the wells were divided into six different groups for further culturing: cells cultured in STD-M (-) for normoxia and hypoxia-derived secretomes, Cyto-M (-) for cytokine-derived secretomes, and HG-M (-) for high glucose and hypoxia + high glucose-derived secretomes. The cells were cultured under either normoxia (21% O_2_, 5% CO_2_) or hypoxia (1% O_2_, 5% CO_2_) conditions for 72 hours. After the 72-hour incubation, the cells’ supernatant (secretomes) from h-prASC and r-prASC was collected and centrifuged (2500 x g, 10 min) to remove unwanted components. Four or five secretome pools were used to perform all functional assays. The pools were created by pooling individual secretomes of three different donors. For instance, for n = 4, the experiment was performed four times with four different pooled secretomes. Pooled secretomes were aliquoted and snap-frozen in liquid nitrogen and stored at −80°C until processing.

### Evaluating the prASC secretomes’ proangiogenic potential

#### Human and rat endothelial cells

To compare the *in vitro* proangiogenic potential of the h-prASC secretomes, Human Umbilical Vein Endothelial Cells (HUVEC) purchased from Lonza Bioscience (Breda, The Netherlands) and cultured by the Endothelial Cell facility of the UMCG (Groningen, The Netherlands) were used. HUVEC were cultured in EBM-2 (+) medium at 37 °C and 5% CO_2_. To compare the *in vitro* proangiogenic potential of the r-prASC secretomes, rat aortic endothelial cells (RAEC) were isolated following a previously described protocol by Suh *et al.* (37). Briefly, the aortas of the same animals previously used for adipose tissue collection were removed and placed in ice-cold PBS. The artery was then carefully cleaned, removing the periadventitial fat and connective tissue. Clean aortas were opened longitudinally and cut into small (2mm) pieces. The pieces were placed with the intima side down in Cultrex BME (R&D systems; Minneapolis, USA) coated wells of a 6-well plate. A small volume (80 uL) of EBM-2 (+) medium was carefully added to the aorta pieces. These were then incubated for 24 h at 37 °C and 5% CO_2._ After 24 h, the pieces were well-attached, and more EBM-2 (+) was added. After 5–7 days, the aortic pieces were removed, and the endothelial cells migrated from the aortic segments were allowed to grow to confluence. Once confluent, the medium was removed, and the cells were incubated for 60 min at 37 °C in dispase II (Gibco) for recovery from the Cultrex BME. After 60 min, EBM-2 (+) was added, and cells were harvested and centrifuged (500 x g, 5 min). The supernatant was removed, and cells were again centrifuged (500 x g, 5 min). The pellet was washed twice with PBS, and cells were then resuspended in EBM-2 (+) for seeding and culturing in a T25 flask (passage 1). The medium was renewed every three days. HUVEC (passage 4) and RAEC (passage 3) were used for the TFA to test the angiogenic potential.

#### TFA

For the TFA, the HUVEC (n = 5) and RAEC (n = 4) (35.000 cells/well in a 96 wells plate) were seeded onto growth factor-reduced Matrigel (Corning) with the different secretomes (final protein concentration of 0.62 ± 0.3 ug/uL) for 18 hours at 37 °C and 5% CO_2_. After the incubation, a Leica MZ7.5 microscope attached to a Leica IC90 E camera was used to image each well. HUVEC and RAEC cultured in STD-M (-) served as control. The angiogenic potential was defined as the number of branching points identified in each one of the conditions after tube formation. Branching points were counted using Fiji (38). The number of branching points was normalized to the control (STD-M (-); set at 1).

### Evaluating the prASC secretomes’ capacity to stimulate collagen deposition

#### Human and rat fibroblasts

To compare the h-prASC *in vitro* potential to stimulate collagen secretion and deposition, commercially derived human fetal lung fibroblasts (FLF92, passage 28 - 30) cultured in STD-M (+) were used. To compare the r-prASC *in vitro* potential to stimulate collagen secretion and deposition, rat dermal fibroblasts (RDF) were used. To obtain those, biopsies were taken from the skin of the same animals used for adipose tissue collection, and a serial explant technique for isolating rat dermal fibroblasts was used as previously described by Nejaddehbashi *et al.* (39). Briefly, skin biopsies (2 x 2 cm^2^) were obtained from the animals, and once collected, the skin samples were kept on ice in STD-M (+) supplemented with gentamicin amphotericin B (0.5 ug/mL; Sigma-Aldrich). Samples were then quickly submerged in 70% ethanol for sterilization and washed with sterile PBS. The surrounding blood vessels and hypodermal adipose tissues were removed, and samples were cut into smaller pieces (3 x 4 mm^2^). The epidermal layer was then digested in 0.3% collagenase I (Sigma-Aldrich) in a water bath at 37 °C for 30 min. Upon digestion, the remaining undigested dermis was placed in one drop of DMEM/F12 in a petri dish (two pieces per dish). Fibroblast outgrowth was observed the next day, and the explants were removed from the Petri dish. The cells were carefully washed with PBS, and medium (DMEM/F12) was added. After three days in culture, the cells were serially cultured into new Petri dishes every time they achieved ∼80% confluence until day 10. The medium was changed every three days. Cultured RDF were used for experiments between passages 12 and 15.

#### Stimulation of collagen deposition

Picrosirius-Red staining (Direct Red 80, Sigma-Aldrich) was used to detect and quantify collagen deposition by FLF92 and RDF cells upon exposure to the different human or rat prASC secretomes. To this end, we adapted the protocol previously described by Xu *et al.* (40). In brief, on the first day, cells were seeded (30,000 cells/well) and cultured in 24-well plates overnight in STD-M (+). The following day, the medium was discarded, cells were carefully washed three times with PBS, and STD-M (-) was added for overnight starvation. On the third day, the medium was removed, and cells were exposed to the different prASC secretomes (human or rat, respectively) for 72 h (n = 4). STD-M (-) is used as control. The medium was not changed during this period.

#### Picrosirius-Red staining

After 72 hours, the cells were fixed in ice-cold methanol (Merck, Darmstadt, Germany) overnight at - 20°C. The following day, cells were carefully washed once with PBS and incubated in the Picrosirius-Red staining solution at RT for 1 h. The staining solution was removed, and the cells were washed three times with 0.1% acetic acid (Merck). Images were taken from each well prior to the spectrophotometric analysis. For that, plates were let to air-dry. Afterward, the Picrosirius-Red dye staining the cells in each of the wells was eluted by adding sodium hydroxide (200 uL/well NaOH, 1M (Merck)) while shaking for 1 h at RT. The eluted dye was collected, and the samples’ optical density (OD) was measured in a 96 wells plate at 540 nm using a spectrophotometer (Epoch 2, BioTek, Winooski, USA). ODs were corrected by the total number of cells counted (Countess 3 Cell Counter, ThermoFisher) upon exposure to each one of the conditions. Collagen content is represented as OD per 10.000 cells. ODs were normalized to the control (STD-M (-); set at 1).

### Evaluating the prASC secretomes’ immunomodulatory potential

To investigate the immunomodulatory potential of the different h-prASC secretomes, we adapted a previously described protocol (41, 42). This protocol includes a one-way MLR followed by an antibody-mediated cell-dependent cytotoxicity (CDC) assay.

#### Peripheral blood mononuclear cells (PBMC)

To test the effect of different human prASC, isolated human PBMC were used. To this end, buffy coats of healthy blood donors were obtained from the Sanquin Blood Bank (Groningen, The Netherlands). A Lymphoprep density gradient (Fresenius Kabin Norge AS, Oslo, Norway) and centrifugation (800 x g, 15 min, RT) were used to obtain the PBMC. Subsequent centrifugation to wash the cells was carried out at 300 x g (5 min, RT). The supernatant was discarded, and the interphase consisting of PBMC was transferred to new centrifuge tubes and washed three times with PBS. Before the final centrifugation, the PBMC were filtered through a 100 µm cell strainer (FALCON, Corning, Durham, USA). The viable cells were counted and immediately used for the MLR (counting and viability assessments were performed using a Countess 3 Cell Counter (ThermoFisher)).

To investigate the immunomodulatory potential of the different r-prASC secretomes, rat splenocytes were used. To obtain those, spleens were collected from the same animals used for adipose tissue collection as previously described (43). Briefly, the spleens were cut into small pieces and mechanically disrupted in ice-cold RPMI (+) (Gibco). Splenic red blood cells were eliminated by incubation with ice-cold ammonium chloride (4 mL, 10 min; UMCG Pharmacy). Falcon tubes with cell strainer caps (35 μm; Corning) were used to remove cell clumps before the cells were counted and immediately plated for the MLR.

#### One-way MLR followed by an antibody-mediated CDC assay

Humoral alloimmunity was evaluated using an adapted method as previously outlined (41, 42). Briefly, the first phase of the assay involved a co-culture between stimulator and responder cells. These were PBMC or splenocytes obtained from mismatched human or rat donors (stimulators: Sprague Dawley rats; responders: Wistar rats). This co-culture was maintained at 37 °C and 5% CO_2_ for seven days in the presence or absence of the different secretomes. During this period, alloantibodies were generated. Simultaneously, resting PBMC or splenocytes (obtained from the same donor as stimulator cells) were cultured for seven days at 37 °C and 5% CO_2_. At the end of the incubation period, the supernatant resulting from the stimulators and responders’ cell interaction was harvested. Also, the resting cells were collected and counted. In the subsequent phase, an antibody-mediated CDC assay was performed. The resting cells were cultured at RT in the supernatants containing alloantibodies for 30 min. Subsequently, a low-toxicity rabbit complement (Sanbio B.V., Cedarlane, Uden, The Netherlands) was introduced, and the culture was continued for 2 h at 37 °C and 5% CO_2_. PBMC or splenocytes’ survival upon interacting with the alloantibodies was assessed using a WST-1 assay (Sigma-Aldrich) according to the manufacturer’s instructions. The efficacy of each secretome in generating alloantibodies with the potential to hamper humoral alloimmunity was evaluated in terms of the percentage of human PBMC or rat splenocytes that survived exposure to these alloantibodies. The survival of cells was normalized to the control (CMRL (-), set at a reference value of 100).

### Statistics

Data are presented as mean ± standard deviation. A Shapiro–Wilk normality test was performed to test the data for normality. The data were statistically analyzed by one-way ANOVA followed by Tukey’s *post hoc* test or by a Kruskal-Wallis test with Dunn’s *post hoc* test. The analyses were performed using GraphPad Prism Software v. 8.3.0 (GraphPad Prism Software, San Diego, CA, USA). Differences were considered significant if p < 0.05.

## Results

### The proangiogenic capacity of prASC secretomes

To determine the capacity of the different prASC secretomes to promote the formation of capillary-like tubes – an indicator of the secretomes’ proangiogenic potential – a TFA was performed. To this end, human (HUVEC) and rat (RAEC) endothelial cells were exposed to the respective h- and r-prASC secretomes for 20 hours, and the number of branching points within the developed tubular network was quantified. A descriptive overview of the statistical results obtained in both human and rat TFA can be found in Supplementary Tables 1 and 2.

#### All h-prASC secretomes stimulate the formation of tube-like structures by HUVEC

Figures 1A-F show representative images of the TFA after incubation of HUVEC with the secretomes derived from different culturing conditions. Figure 1G shows an analysis of the branching points compared among all tested conditions. The results showed that HUVEC exposed to the h-prASC normoxia (Fig. 1B) or hypoxia-derived (Fig. 1E) secretomes formed the highest number of branching points (Fig. 1G). The number of branching points resulting from HUVEC exposed to these secretomes was significantly higher when compared to HUVEC exposed only to STD-M (-) (Fig. 1A). A 2.24-fold-change increase in branching points was observed when HUVEC were exposed to the h-prASC normoxia-derived secretome (p < 0.0001 vs. STD-M (-)), and more robust increase (2.44-fold-change) was detected in HUVEC exposed to the h-prASC hypoxia-derived secretome (p < 0.005 vs. STD-M (-)). Exposure to the cytokines (Fig. 1C), high glucose (Fig. 1D), and hypoxia + high glucose-derived (Fig. 1F) secretomes also significantly impacted the number of branching points formed by HUVEC compared to the control (Fig. 1G). Respectively, a 1.59, 1.78, and 1.74-fold-change increase (vs. STD-M (-)) was detected.

**Figure 1.**
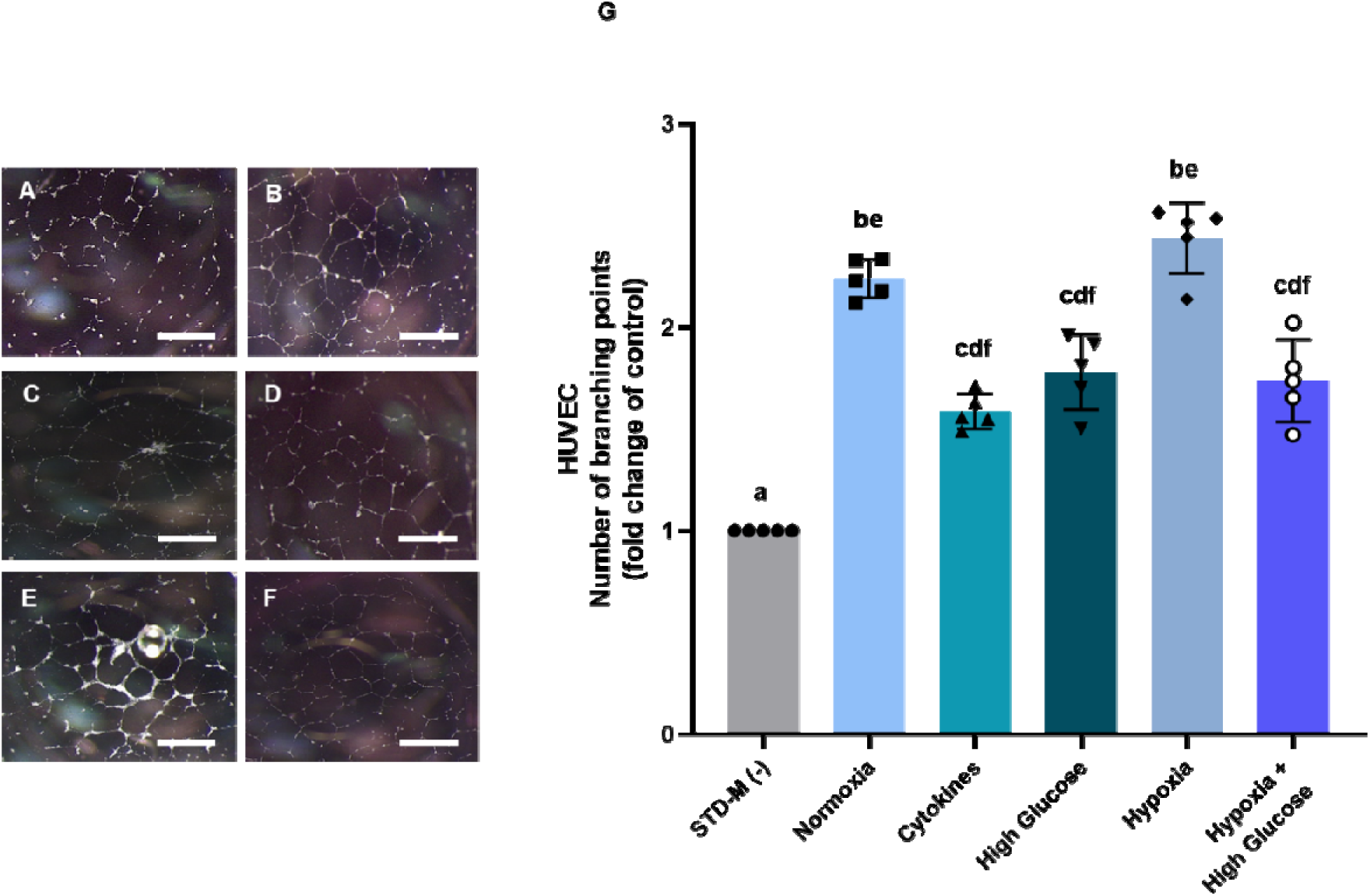
Human prASC secretomes enhance HUVEC angiogenic capacity. (A-F) Representative images of tube formation with HUVEC exposed to the various h-prASC secretomes for 20 hours. Scale bar, 100 µm. Exposure to (A) STD-M (-); (B) h-prASC normoxia secretome; (C) h-prASC cytokines secretome; (D) h-prASC high glucose secretome; (E) h-prASC hypoxia secretome; (F) h-prASC hypoxia + high glucose secretome. (G) Quantification of the branching points formed upon exposure (n = 5). The number of branching points was normalized to the control (STD-M (-); set at 1). Data are represented as individual values and mean ± SD. Statistical analysis was carried out using a One-way ANOVA with Tukey’s *post hoc* test. Different letters indicate significant differences, p < 0.05. Detailed description of the differences can be found in Supplementary table 1.

When comparing the proangiogenic capacity among the secretomes, no differences were detected between the normoxia and hypoxia-derived h-prASC secretomes’ capacity to stimulate HUVEC tube formation (Fig. 1G). Specifically, the normoxia-derived secretome exhibited a 1.41-fold increase over cytokines (p < 0.005), a 1.25-fold increase over high glucose (p < 0.05), and a 1.28-fold increase over hypoxia + high glucose-derived secretomes (p < 0.05), respectively. On the other hand, the hypoxia-derived secretome demonstrated a 1.54-fold increase over cytokines, a 1.37-fold increase over high glucose (both p < 0.001 compared to hypoxia), and a 1.40-fold increase over hypoxia + high glucose (p < 0.05 compared to hypoxia). Moreover, no differences were observed when comparing the capacity to stimulate tube formation among the cytokines, high glucose, and hypoxia + high glucose h-prASC secretomes. For the latest, when considering hypoxia and high glucose as a culturing condition, their combined effect did not result in a secretome that further stimulated HUVEC tube formation (Fig. 1G) compared to using hypoxia or high glucose individually. HUVEC exposed to the h-prASC hypoxia-derived secretome formed 1.40 times more branching points than HUVEC exposed to the h-prASC hypoxia + high glucose secretome.

#### All r-prASC secretomes except for the high glucose-derived one stimulate the formation of tube-like structures by RAEC

Figures 2A-F show representative images of the TFA after incubation of RAEC with the secretomes derived from different culturing conditions. Figure 2G shows the analysis of the branching points compared among all conditions tested. The data showed that, compared to the control (STD-M (-)), all r-prASC secretomes increased the number of branching points formed by RAECS except for the r-prASC high glucose secretome (Fig. 2G). The highest significative responses were detected in RAEC exposed to the r-prASC normoxia (Fig. 2B), cytokines (Fig. 2C), and hypoxia-derived (Fig. 2E) secretomes. Respectively, the number of branching points was 1.940, 1.28, and 2.39 times higher than the control (p < 0.01 for all). To a lesser extent, RAEC exposed to the hypoxia + high glucose-derived r-prASC secretome (Fig. 2F) also displayed an increased number of branching points (p < 0.05) when compared to the control (Fig. 2G).

**Figure 2.**
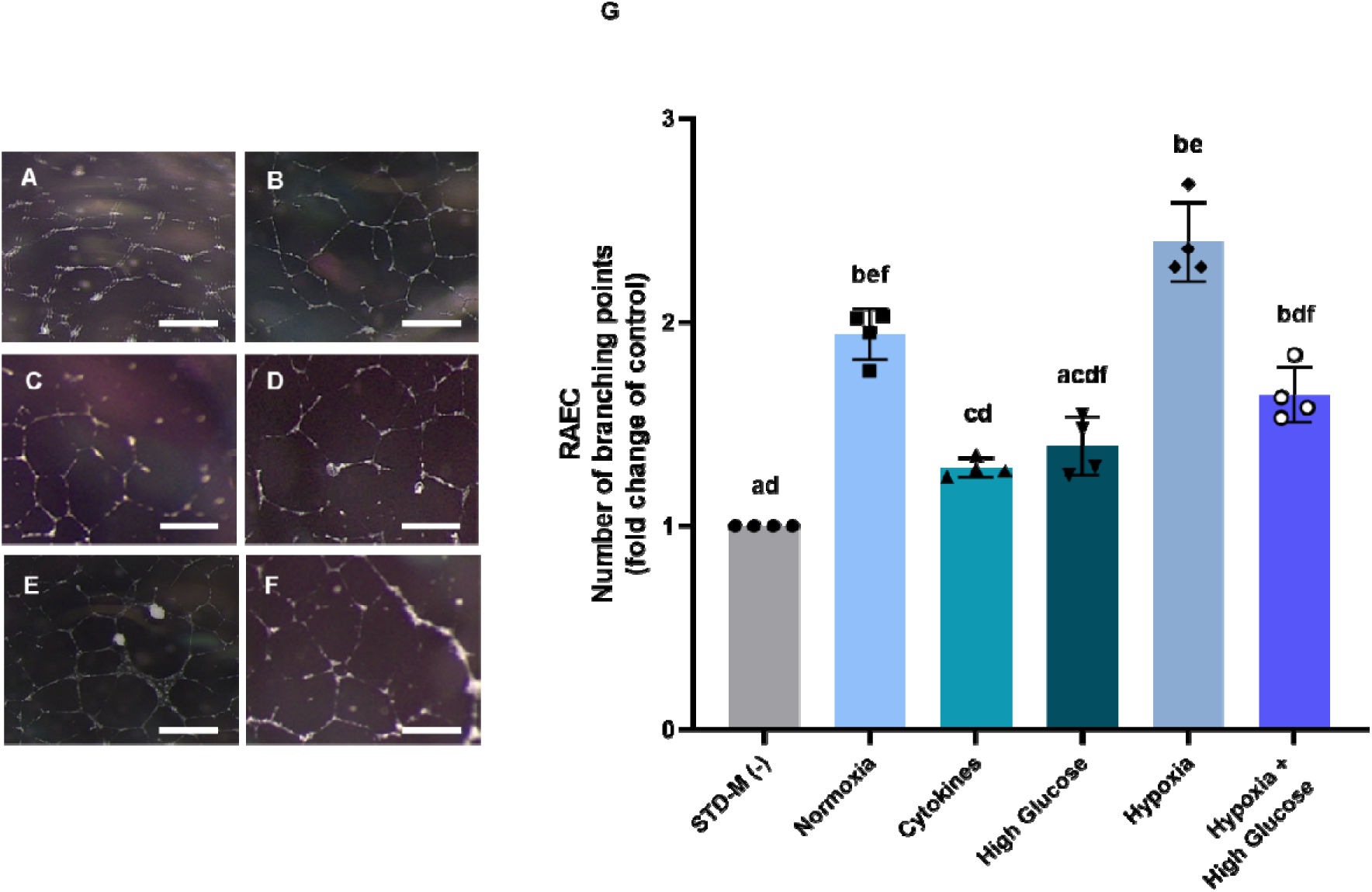
Rat prASC secretomes enhance RAEC angiogenic capacity. (A-F) Representative images of tube formation with RAEC exposed to the various r-prASC secretomes for 20 hours. Scale bar, 100 µm. Exposure to (A) STD-M (-); (B) r-prASC normoxic secretome; (C) r-prASC cytokines secretome; (D) r-prASC high glucose secretome; (E) r-prASC hypoxia secretome; (F) r-prASC hypoxia + high glucose secretome. (G) Quantification of the branching points formed upon exposure (n = 4). The number of branching points was normalized to the control (STD-M (-); set at 1). Data are represented as individual values and mean ± SD. Statistical analysis was carried out using a one-way ANOVA with Tukey’s *post hoc* test. Different letters indicate significant differences, p < 0.05. Detailed description of the differences can be found in Supplementary table 2.

Comparisons among the secretomes revealed that the normoxia-derived secretome was a more potent stimulator for RAEC tube formation than the cytokines and high glucose-derived secretomes (p < 0.05; 1.51 and 1.40 times increase) but as potent as the hypoxia and hypoxia + high glucose-derived secretomes (Fig. 2G). The cytokines and high glucose-derived secretomes were equally capable of stimulating RAEC formation, with a decreased capacity to stimulate tube formation compared to the hypoxia-derived secretome (p < 0.001 cytokines; p < 0.05 high glucose vs. hypoxia). Compared to the high glucose-derived secretome, no additive stimulus was observed when RAEC were exposed to the hypoxia + high glucose-derived secretome. The latter also resulted in significantly fewer branching points than the hypoxia-derived secretome (p < 0.05).

### The capacity of the prASC secretome to stimulate collagen deposition

To evaluate the effect of the prASC secretomes on the deposition of ECM, the production of collagen, the most abundant ECM component, was analyzed using Picrosirius-Red staining and spectrometric analysis. In this way, investigating how h- and r-prASC secretomes derived from various culturing conditions could affect the capacity of human and rat fibroblasts to secrete collagen after 72 hours of exposure. A descriptive overview of the statistical results obtained in both human and rat Picrosirius-Red staining assays can be found in Supplementary Tables 3 and 4.

#### Normoxia, hypoxia, and hypoxia + high glucose-derived h-prASC secretomes stimulate collagen deposition by human fibroblasts

Figures 3A-F show representative images of the Picrosirius-Red staining on human fibroblast exposed to the h-prASC secretomes derived from different culturing conditions. Figure 3G shows the OD values of the recovered dye per 10.000 cells per condition. The data showed that, compared to the control (STD-M (-)), a stimulative effect on collagen deposition was observed upon fibroblasts’ exposure to the normoxia (Fig. 3B; p < 0.001), hypoxia (Fig. 3E; p < 0.001), and hypoxia + high glucose-derived (Fig. 3F; p < 0.05) secretomes (Fig. 3G). Respectively, we detected an increase of 1.47, 1.64, and 1.30 times in the OD upon the Picrosirius-Red dye recovery in each one of these conditions. The high glucose-derived secretome (Fig. 3D) was the only h-prASC secretome that induced a significative negative effect on the amount of collagen deposited by fibroblasts – a decrease of 0.7 times (p < 0.05 vs. STD-M (-)).

**Figure 3.**
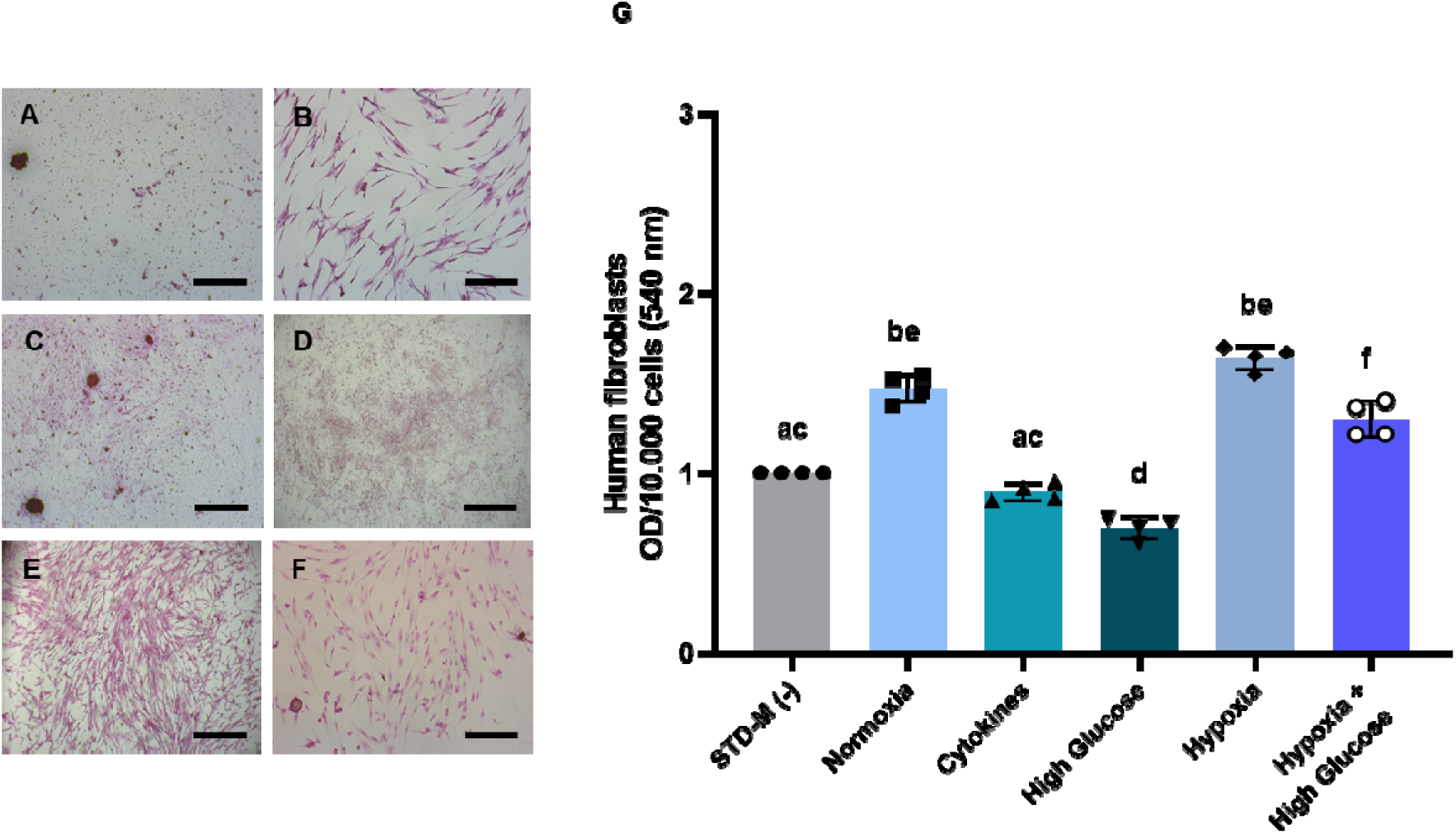
Human prASC secretomes stimulate collagen secretion by human fibroblasts. (A-F) Representative images of Sirius-Red staining in human fibroblasts exposed to various h-prASC secretomes for 72 hours. Scale bar, 50 µm. Exposure to (A) STD-M (-); (B) h-prASC normoxic secretome; (C) h-prASC cytokines secretome; (D) h-prASC high glucose secretome; (E) h-prASC hypoxia secretome; (F) h-prASC hypoxia + high glucose secretome. (G) Spectrophotometric analysis of the Sirius-Red-stained human fibroblasts depicted in A (n = 4). Data are represented as individual values and mean ± SD. Statistical significance was assessed using one-way ANOVA and Tukey’s *post hoc* test. Different letters indicate significant differences, p < 0.05. Detailed description of the differences can be found in Supplementary table 3.

When comparing the capacity to stimulate fibroblasts’ collagen deposition among all h-prASC secretomes, no differences were detected between fibroblasts exposed to normoxia and hypoxia-derived secretomes (Fig. 3G). Respectively, the OD values resulting from exposure to these conditions were 1.63 and 1.82 times higher (p < 0.001) than fibroblasts exposed to the cytokines-derived secretome. They were also, respectively, 2.10 (p <0.005) and 2.34 (p < 0.0001) times higher than fibroblasts exposed to the high glucose-derived secretome. A potential additive effect of the hypoxia + high glucose secretome was not observed here. Fibroblast exposure to this secretome resulted in OD values that were 1.85 times higher than the ones detected upon fibroblast exposure to the high glucose-derived secretome (p < 0.001) but 0.8 times lower (p < 0.05) than the ones detected in cells exposed to the hypoxia-derived secretome.

Differences were also observed regarding the spatial distribution and morphology of the collagen deposits stained with Picrosirius-Red. Spatially, the collagen fibers deposited by fibroblasts exposed to normoxia, hypoxia, and hypoxia + high glucose-derived secretomes seemed to display a more linear orientation (Fig. 3B, E, F), with collagen in bundles. In contrast, fibroblasts exposed to the cytokines and high glucose-derived secretomes displayed more diffuse staining (Fig. 3C-D), in which denser, rounded-like structures can be observed as in collagen nodules (Fig. 3C).

#### Normoxia and hypoxia-derived r-prASC secretomes stimulate collagen deposition by rat fibroblasts

Figures 4A-F show representative images of the Picrosirius-Red staining on rat fibroblasts exposed to the r-prASC secretomes derived from different culturing conditions. Figure 4G shows the OD values of the recovered dye per 10.000 cells per condition. The data showed that, compared to the control (STD-M (-), Fig. 4A), rat fibroblasts exposed to the normoxia (Fig. 4B) and hypoxia-derived (Fig. 4E) secretomes presented a significantly higher amount of collagen deposition (Fig. 4G). Respectively, a 1.68 (p < 0.05) and 2.18 times (p < 0.05) increase in the OD of the Sirius-Red staining was found for these groups. No significant differences were found in the collagen deposition of cells exposed to the cytokines (Fig. 4C), high glucose (Fig. 4D), and hypoxia + high glucose secretomes (Fig. 4E) as compared with the control (STD-M (-)).

**Figure 4.**
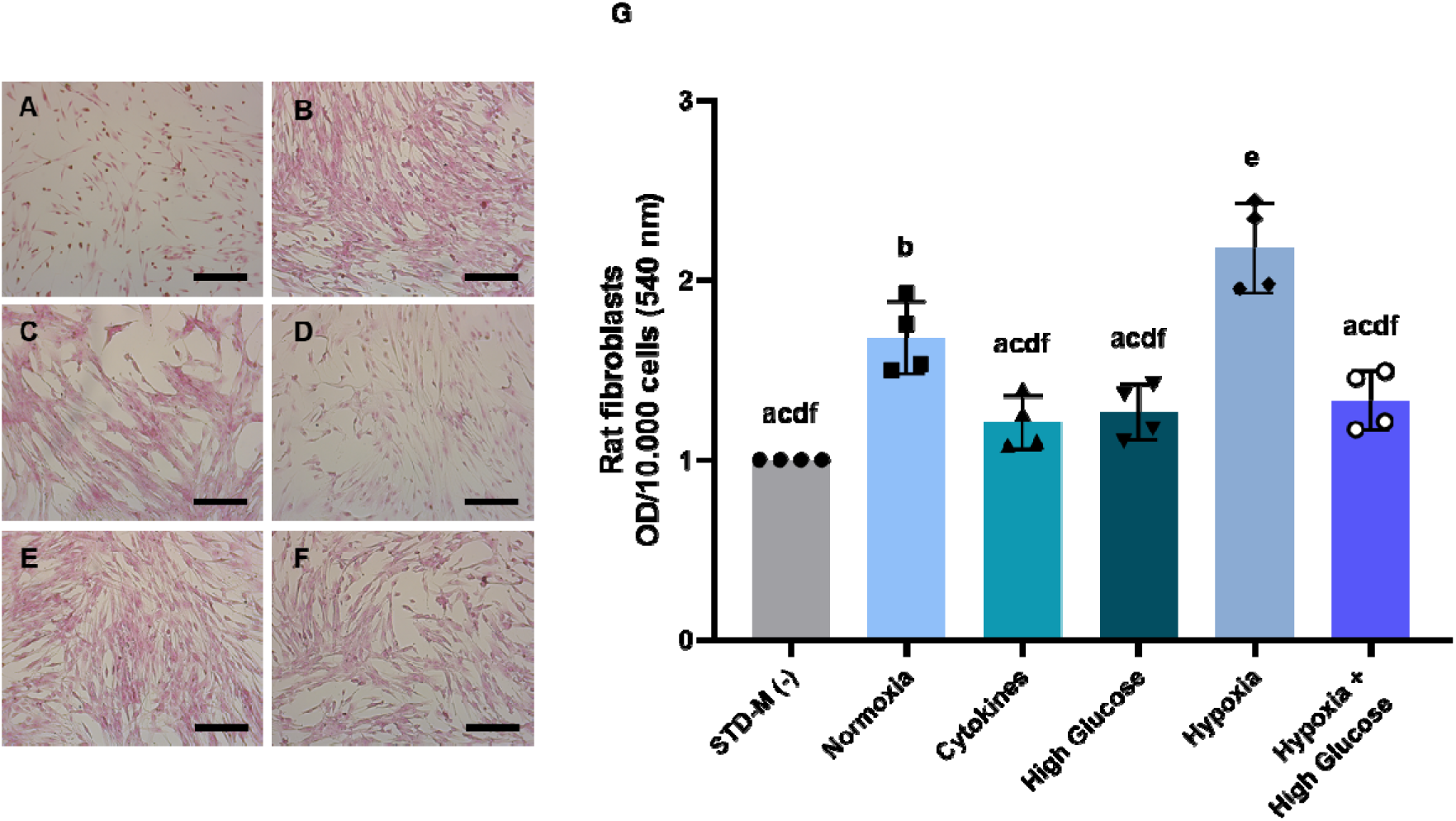
Rat prASC secretomes stimulate collagen secretion by rat fibroblasts. (A) (A-F) Representative images of Sirius-Red staining in rat fibroblasts exposed to various r-prASC secretomes for 72 hours. Scale bar, 50 µm. Exposure to (A) STD-M (-); (B) r-prASC normoxic secretome; (C) r-prASC cytokines secretome; (D) r-prASC high glucose secretome; (E) r-prASC hypoxia secretome; (F) r-prASC hypoxia + high glucose secretome. (G) Spectrophotometric analysis of the Sirius-Red-stained rat fibroblasts depicted in A (n = 4). Data are represented as individual values and mean ± SD. Statistical significance was assessed using one-way ANOVA and Tukey’s post-hoc test for multiple comparisons correction. Different letters indicate significant differences, p < 0.05. Detailed description of the differences can be found in Supplementary table 4.

When comparing the secretomes among each other, the hypoxia-derived secretome (Fig. 4E) showed the most significant stimulatory effect on collagen deposition of all r-prASC secretomes tested (Fig. 4G, p < 0.001 vs. normoxia, cytokines, high glucose, and hypoxia + high glucose). This effect was followed by the normoxia-derived secretome, which stimulated 1.38, 1.33, and 1.26 times more deposition of collagen by fibroblasts when compared to, respectively, the cytokines (p < 0.005), high glucose (p < 0.005), and hypoxia + high glucose-derived (p < 0.005) r-prASC secretomes. A potential additive effect of hypoxia + high glucose as a culturing condition for generating a secretome able to enhance fibroblast collagen deposition was also not observed here. OD values resulting from the stained cells exposed to the hypoxia + high glucose-derived secretome were comparable to the ones obtained from cells exposed to the high glucose-derived secretome but less than the hypoxia-derived secretome (p < 0.005).

Unlike what was observed for human fibroblasts exposed to h-prASC secretomes, spatial and morphological differences were not evident here. The collagen staining patterns from rat fibroblasts exposed to all conditions showed the predominance of collagen bundles and scarcity in collagen nodules (Fig 4A – F).

### The immunomodulatory capacity of prASC secretome

The capacity to influence humoral alloimmunity is an important tool in evaluating the immunomodulatory potential of a compound. Humoral alloimmunity refers to an immune response mediated by antibodies against antigens from a foreign body (41). For instance, this response occurs when an individual’s immune system recognizes and responds to non-self-antigens – as in an organ transplantation context (41). To evaluate the prASC secretomes’ immunomodulatory capacity, humoral alloimmunity was assessed using an MLR followed by an antibody-mediated CDC. To this end, the survival of human PBMC and rat splenocytes was measured upon exposure to alloantibodies resulting from the co-culture of cells from unmatched donors in the presence or absence of h- or r-prASC secretomes. A descriptive overview of the statistical results obtained in both human and rat MLR can be found in Supplementary Tables 5 and 6.

#### All h-prASC secretomes, except for the high glucose-derived secretome, increase PBMC survival in an antibody-mediated CDC assay

Figure 5 illustrates the survival percentage of human PBMC after the antibody-mediated CDC assay. The efficiency of this assay was evidenced by a significant decline (p < 0.0001) in PBMC survival when comparing resting PBMC exposed solely to STD-M (+) and those exposed to the MLR supernatant, which did not contain any secretome. Furthermore, when compared to all MLR supernatants resulting from secretome exposure, STD-M (+) exhibited a significantly higher capacity to support PBMC survival (p < 0.0001 DMEM (-) vs. all).

**Figure 5.**
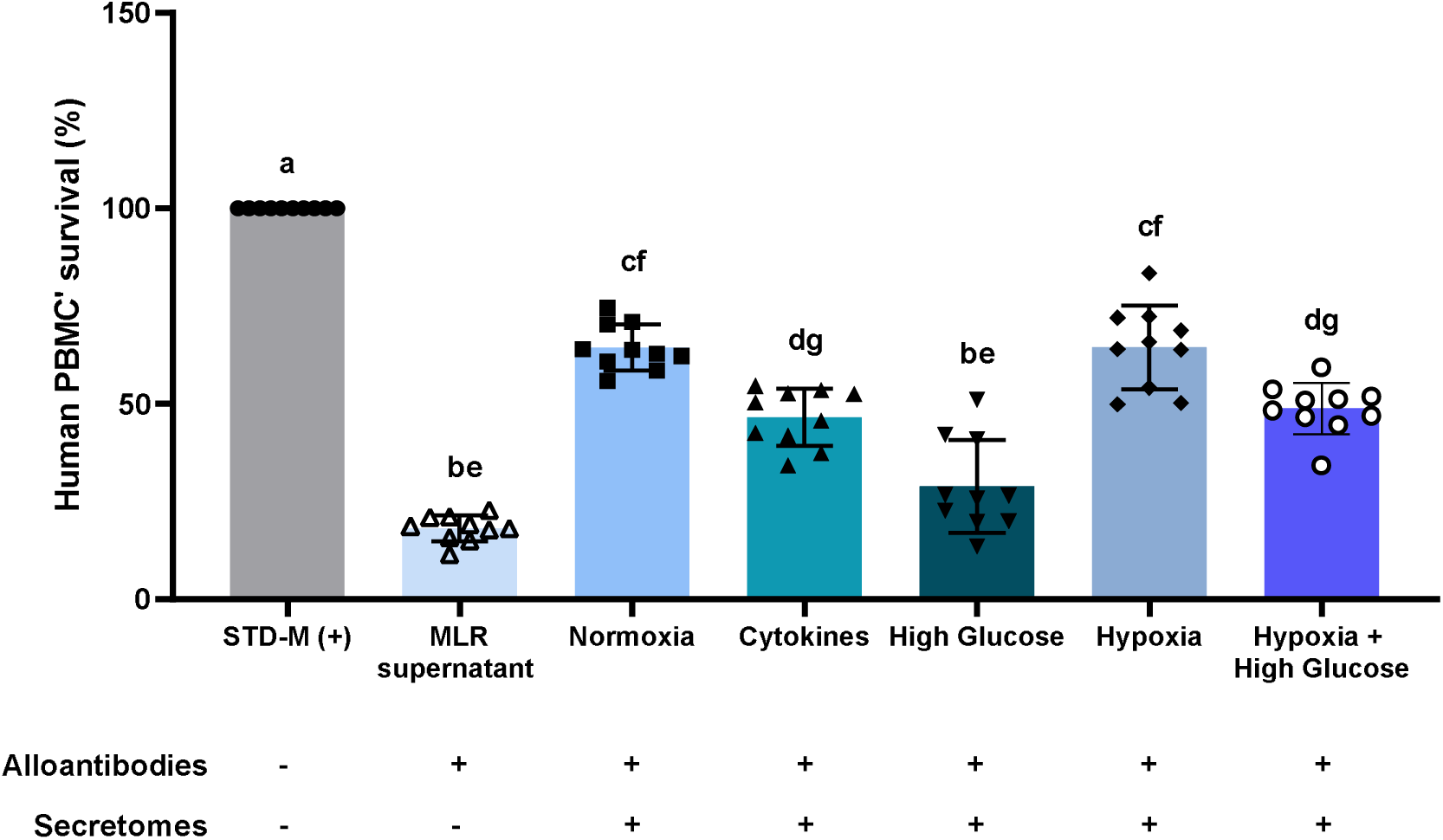
Human prASC secretomes modulate antibody-mediated immune responses. The effect of various h-prASC secretomes on humoral alloimmunity was evaluated using a mixed lymphocyte reaction (MLR) followed by an antibody-mediated complement-dependent cytotoxicity (CDC) assay. The MLR consisted of generating alloantibodies by unmatched human PBMC in the presence or absence of the various h-prASC secretomes during 7-days of culturing. The MLR supernatant contained alloantibodies and was used in an antibody-mediated complement-derived cytotoxicity assay. The outcome of this assay is expressed as the percentage of PBMC that survived the exposure to the alloantibodies (n = 10). RPMI (-) was used as a control and together with the MLR supernatant it shows the efficacy of the antibody-mediated CDC assay. Cell survival was normalized to the control (STD-M (+); set to 100). Data are represented as mean ± SD. Statistical significance was assessed using one-way ANOVA and Tukey’s *post hoc* test. Different letters indicate significant differences, p < 0.05. Detailed description of the differences can be found in Supplementary table 5.

The data revealed an increase in the survival of resting PBMC exposed to MLR supernatants generated in the presence of all human secretomes (p < 0.0001 for normoxia, cytokines, hypoxia, and hypoxia + high glucose vs. MLR supernatant), with the exception of the high glucose-derived secretome, which exhibited no significant effect on survival compared to PBMC exposed to the alloantibodies only (MLR supernatant; Fig. 5). The survival rates increased by 254.9, 156.6, 255.2, 168.8% for normoxia, cytokines, hypoxia and hypoxia + high glucose secretomes, respectively (Fig. 5).

Within the different secretomes-derived MLR supernatants, the highest survival rates were observed upon exposure to the MLR supernatant obtained in the presence of the h-prASC normoxia-derived secretome (p < 0.0001 vs. cytokines and high glucose; p > 0.005 vs. hypoxia + high glucose) or to the hypoxia-derived secretome (p < 0.0001 vs. cytokines and high glucose; p < 0.005 vs. hypoxia + high glucose), suggesting that these secretomes are both highly capable of limiting alloantibodies production in a one-way MLR using human PBMC when compared to all the other secretomes tested. Exposure to the cytokines-derived MLR supernatant resulted in higher PBMC survival compared to the high glucose-derived MLR supernatant (p < 0.005) but equally maintained survival when compared to the hypoxia + high glucose-derived MLR supernatant.

#### All r-prASC secretomes increase splenocytes’ survival in an antibody-mediated CDC assay

Figure 6 shows the survival percentage of rat splenocytes after the antibody-mediated CDC assay. The efficiency of the antibody-mediated CDC assay was demonstrated by a significant decline (p < 0.0001) in splenocytes’ survival when comparing resting splenocytes exposed to STD-M (+) and those exposed to the MLR supernatant, which did not contain any secretomes. Additionally, when compared to all MLR supernatants resulting from secretome exposure, STD-M (+) exhibited a significantly higher capacity to support PBMC survival (p < 0.0001 STD (+) vs. all).

**Figure 6.**
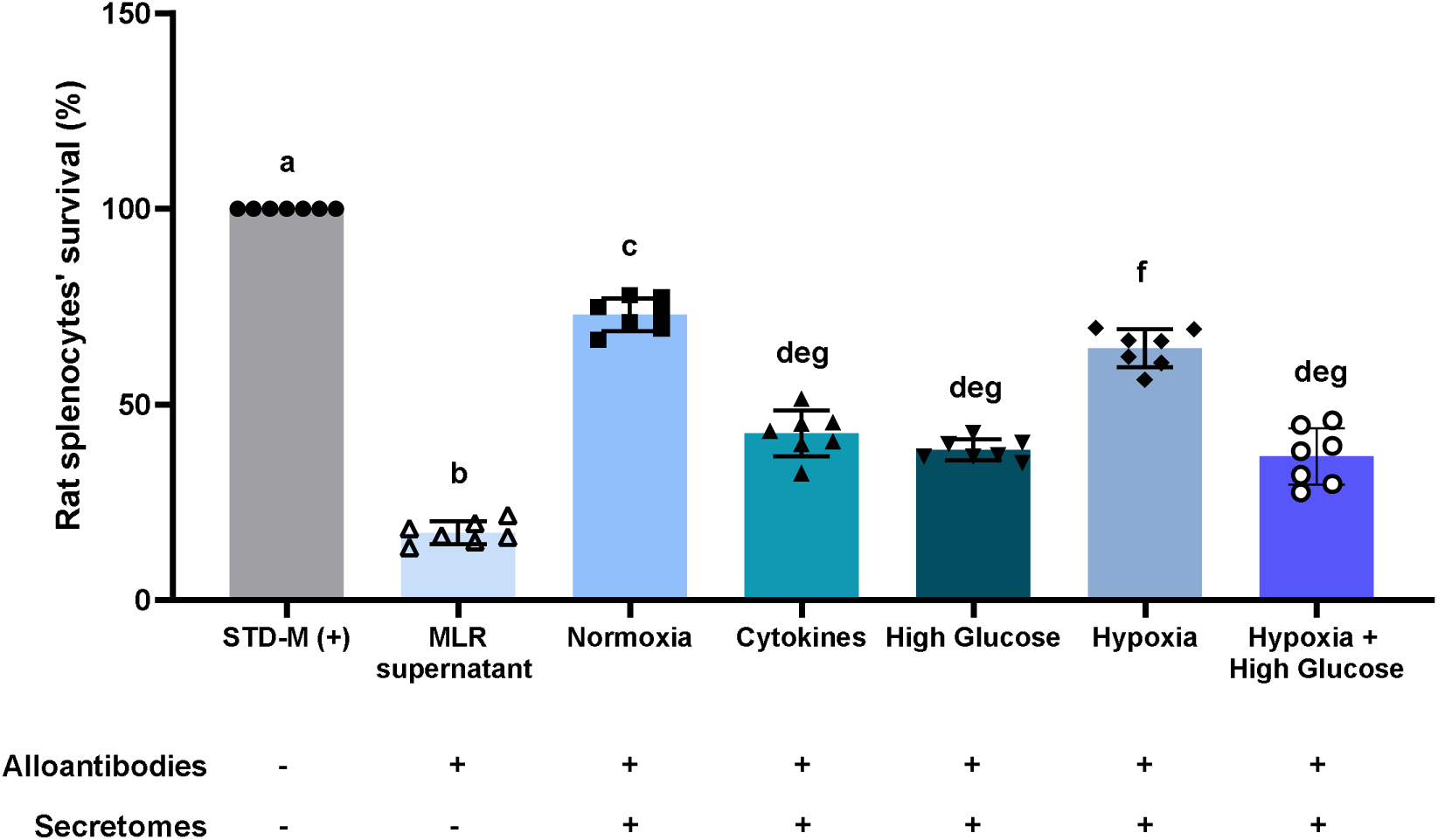
Rat prASC secretomes modulate antibody-mediated immune responses. The effect of various r-prASC secretomes on humoral alloimmunity was evaluated using a mixed lymphocyte reaction (MLR) followed by an antibody-mediated complement-dependent cytotoxicity (CDC) assay. The MLR consisted of generating alloantibodies by unmatched rat splenocytes in the presence or absence of the various r-prASC secretomes during 7-days of culturing. The MLR supernatant contained alloantibodies and was used in an antibody-mediated complement-derived cytotoxicity assay. The outcome of this assay is expressed as the percentage of splenocytes that survived the exposure to the alloantibodies (n = 7). RPMI (-) was used as a control and together with the MLR supernatant it shows the efficacy of the antibody-mediated CDC assay. Cell survival was normalized to the control (RPMI (-); set to 100). Data are represented as mean ± SD. Statistical significance was assessed using one-way ANOVA and Tukey’s *post hoc* test. Different letters indicate significant differences, p < 0.05. Detailed description of the differences can be found in Supplementary table 6.

The data revealed an increase in the survival of rat splenocytes exposed to MLR supernatants generated in the presence of all rat secretomes (p < 0.0001 normoxia and hypoxia; p < 0.001 high glucose; p < 0.01 cytokines; p < 0.05 hypoxia + high glucose vs. MLR supernatant) (Fig. 6). Increases of 325.5, 149.1, 123.4, 274.9, and 114.9% we identified for normoxia, cytokines, high glucose, hypoxia, and hypoxia + high glucose-derived MLR supernatants, respectively.

Within the different secretomes-derived MLR supernatants, the highest survival rate was found in splenocytes exposed to the MLR supernatant obtained in the presence of the normoxia-derived r-prASC secretome (p < 0.0001 vs. high glucose, p < 0.001 vs. cytokines and hypoxia + high glucose; and p < 0.001 vs. hypoxia). Such ability supporting splenocytes’ survival upon exposure to alloantibodies was followed by the MLR supernatant obtained in the presence of the r-prASC hypoxia-derived secretome (p < 0.0001 vs. high glucose, p < 0.001 vs. hypoxia + high glucose, and p < 0.01 vs. cytokines). Splenocytes exposed to MLR supernatants obtained in the presence of the cytokines, high glucose, and hypoxia + high glucose-derived r-prASC secretomes showed no significative change among each other.

## Discussion

Over the years, research has shown the significant contribution of ASC to cell-based therapy. The secretome of these cells, a complex array of bioactive molecules, plays a key role in ASC therapies. Due to the adaptable nature of secretomes, which can be tailored based on ASC culturing strategies and provide a cell-free therapeutic approach, this study has investigated *in vitro* functional implications of distinct prASC secretomes. These include the prASC secretome potential to stimulate angiogenesis, facilitate collagen deposition, and modulate humoral alloimmunity. Together, these investigations shed light on the pivotal role of ASC cell-free therapy in establishing an optimal regenerative environment.

The described findings showed substantial proangiogenic potential from h- and r-prASC secretomes. Among these, normoxia and hypoxia-derived secretomes exhibited the highest proangiogenic effects regardless of species. Angiogenesis is crucial for tissue regeneration, supplying oxygen, nutrients, and immune cells while facilitating growth factor exchange and tissue remodeling (44). Normoxia and hypoxia have been conventionally used as culturing conditions that enhance the ASC secretomes’ angiogenic factors content (10, 28, 45, 46). Previous data, both from our group and others, has consistently demonstrated the enrichment of proteins like VEGF (vascular endothelial cell growth factor), FGF (fibroblast growth factor), and HGF (hepatocyte growth factor) within such secretomes (19, 46, 47). These proteins are known as principal mediators underlying the proangiogenic properties of these ASC secretomes. These factors promote endothelial cell proliferation, migration, and capillary tube formation, thereby facilitating the process of angiogenesis (9, 25, 45, 48). Moreover, our previous data has shown that these secretomes are also enriched in angiogenesis-related pathways such as the VEGFA-VEGFR2 signaling pathway, the regulation of insulin-like growth factors (IGF) transport, and uptake by insulin-like growth factor binding proteins (IGFBPs) (36).

Moreover, our findings demonstrated the ability of cytokines and high glucose-derived prASC secretomes to stimulate tube formation. However, their effectiveness was not as pronounced as that observed with the normoxia and hypoxia-derived counterparts. The cytokines-derived ASC secretome has been mostly reported as immunomodulatory (49, 50) but also has proangiogenic properties (51). The presence of proteins such as VEGF, FGF, IL-6 (interleukin 6), IL-8 (interleukin 8), CCL2 and 7 (monocyte chemoattractant protein 1 and C-C motif chemokine 7), CXCL9 (C-X-C motif chemokine 9), and ANGPT-1 (angiopoietin-1) – crucial players in the angiogenic process – support such capacity (44). Moreover, enrichment in other proteins such as TSG-6 (tumor necrosis factor-inducible gene 6 protein) and IDO (indoleamine 2,3-dioxygenase) suggests a secretome that can regulate ROS (reactive oxygen species) production, influence ECM organization, and promote immune suppression – processes indirectly enhancing angiogenesis (52–54).

In turn, even though there have been no prior investigations into the impact of high glucose on the h-prASC secretion composition or functional profile, our earlier research (36) demonstrated that the secretome of high glucose-derived h-prASC contains proangiogenic proteins. We also showed that this secretome is enriched in immune system pathways, ECM organization, and metabolism – due to the presence of complement proteins, collagens, and glycolytic enzymes (36). This result correlates with our current TFA outcomes, as such pathways are known to play a complex role in angiogenesis. To note, two types of endothelial cells were used in this study – a human vein-derived EC and a rat artery-derived EC. Veins generally have a higher capacity for sprouting new blood vessels than arteries (55), which may partly explain the overall differences we observed in this study between human and rat prASC proangiogenic potential. Regardless of the type of blood vessel used, the secretomes from both species showed significative stimulatory capacity towards forming tube-like structures.

This study also shows the efficiency of normoxia and hypoxia-derived secretomes in stimulating collagen deposition by fibroblasts. In the case of the h-prASC normoxia-derived secretome, it exhibited a similar stimulatory capacity as the h-prASC hypoxic secretome. However, this equivalence was not observed with the r-prASC secretomes, where the r-prASC hypoxic secretome outperformed the r-prASC normoxic secretome in inducing collagen deposition by rat fibroblasts. Such an effect could be attributed to higher amounts of HIF-1 α (hypoxia-induced factor 1α) in the hypoxic-derived secretomes (36). Indeed, our previous findings have established the presence of HIF-1α in both human and rat prASC hypoxia (and hypoxia + high glucose)-derived secretomes (36). HIF-1α is mainly found intracellularly, but it has also been shown to be present in extracellular vesicles (56, 57), which makes this protein part of some cells’ secretome. Under hypoxic conditions, ASC upregulate HIF-1α, a transcription factor (58), which has been shown to promote collagen synthesis and deposition (59, 60). HIF-1α stabilizes the mRNA of collagen molecules by interacting with specific regulatory elements within their mRNA sequences (60, 61). This stabilization effect increases collagen mRNA levels, facilitating enhanced collagen deposition. Additionally, proteins known to stimulate collagen deposition, such as collagen I alpha 2 and collagen I alpha 1-derived proteins, have also been identified in such secretomes (36). These factors potentially contribute to the positive impact of normoxia and hypoxia-derived secretomes on collagen deposition.

More species-dependent variations were identified in terms of collagen deposition when using high glucose-derived prASC secretomes from humans and rats. Specifically, the h-prASC high glucose-derived secretome led to a reduction in collagen deposition by human fibroblasts. Conversely, no discernible impact was observed when using r-prASC high glucose-derived secretomes and assessing collagen deposition by rat fibroblasts. This contrast could potentially arise from the distinct pathways reported to be enriched within these species-dependent prASC high glucose-derived secretomes and the utilization of varying fibroblast types for this study. The r-prASC secretome was enriched in proteins part of the proteasome degradation pathway, while its human counterpart was not (36). This pathway regulates collagen turnover (62, 63), which may ultimately contribute to the improved collagen deposition observed in the rat fibroblasts exposed to the r-prASC high glucose-derived secretome. Additionally, we have employed dermal fibroblasts to evaluate the r-prASC secretomes. Dermal fibroblasts may be less sensitive than lung fibroblasts – used for testing the h-prASC secretomes – as in physiological conditions, dermal fibroblasts are constantly exposed to more dynamic and diverse external stimuli (64, 65). As the h-prASC high glucose-derived secretome was previously described to contain proteins associated with cell programmed death and response to stress pathways (36), it might be possible that the lung fibroblasts were less resilient to these proteins than the dermal cells. Nonetheless, except for the human high glucose-derived prASC secretome, all other secretomes were as much or more capable of positively modulating ECM organization than the controls, which is positive for creating a favorable environment for tissue regeneration.

This study also demonstrated that human and rat prASC secretomes can modulate humoral alloimmunity. This capacity corroborates several previously reported studies describing the immunomodulatory effects of ASC secretomes (5, 27, 54). When using the h-prASC secretomes, it was observed that all secretome types seemed to be able to decrease alloantibody production, as indicated by increased survival of PBMC in the CDC assay, except for the high glucose-derived secretome. Our previous data showed that this secretome was enriched in proteins that participated in protein processing and proteasomal system pathways (36). These include various proteases and ubiquitins implicated in proteolytic degradation processes that may enhance the availability of peptide fragments presented to B cells – increasing alloantibodies production in the MLR context and thus decreasing the viability of PBMC. Furthermore, the h-prASC normoxia and hypoxia-derived secretomes showed the largest effect on PBMC survival.

R-prASC secretomes also showed a high capacity to limit *in vitro* alloantibodies production. Within the antibody-mediated CDC assay using the supernatants resulting from the one-way MLR, the survival percentage of the rat splenocytes was the highest, with the presence of the MLR supernatant obtained in the presence of the normoxia-derived secretome. This indicates that this secretome is the most potent rat secretome, inhibiting the production of alloantibodies. We showed that the presence of MLR supernatant resulting from normoxia and hypoxia-derived h-prASC secretomes showed an equal ability to support human PBMC survival. Here, the MLR supernatant resulting from the r-prASC hypoxia-derived secretome presence displayed a reduced capacity when compared to the one in the presence of the r-prASC normoxia-derived secretome. This could be explained by the reduced enrichment in immune system-related proteins in the r-prASC hypoxia-derived secretome when compared to the h-prASC. We have previously described the r-prASC normoxia secretome as a secretome enriched in pathways associated with oxidative stress and regulation of apoptosis (36).

MLR supernatants resulting from one-way MLR in the presence of the r-prASC cytokines, high glucose, and hypoxia + high glucose-derived secretomes showed reduced support for splenocytes’ survival. We previously reported the presence of the tryptophan metabolism pathway in all of them (36). The effect of tryptophan metabolism on immunity is complex as the pathway plays a role in regulating T cell function and inflammation, energy homeostasis, and many other processes (66). Metabolites, enzymes, and other proteins involved in this pathway are known to contribute to T-cell function and inflammation (66–68). T cells and inflammatory factors are known to affect B cell activation and antibody production, and thus, the presence of factors associated with tryptophan metabolism may explain why these secretomes do not display an enhanced capacity to mitigate alloimmunity in the context explored by us when compared to the r-prASC normoxia and hypoxia-derived secretomes.

## Conclusion

Angiogenesis, collagen deposition, and immunomodulation are intertwined processes crucial for effective tissue regeneration. In conclusion, this study elucidates the distinct effects of prASC secretomes on these processes. Normoxia and hypoxia-derived prASC secretomes consistently enhanced angiogenesis, collagen deposition, and immunomodulation, suggesting their potential as primary candidates for ASC secretome-based regenerative strategies.

## Research data

All the data used in this manuscript is available as a Supplementary Material. Further questions regarding data should be forwarded to the corresponding author: Alexandra M. Smink, PhD..

## CRediT authorship contribution statement

Erika Pinheiro-Machado: Conceptualization, Methodology, Validation, Formal analysis, Investigation, Data curation, Writing – Original Draft, Writing – Review & Editing, Visualization. Claudia C. Koster: Methodology, Investigation, Writing - Review & Editing. Alexandra M. Smink: Conceptualization, Writing – Original Draft, Writing – Review & Editing, Visualization, Supervision, Project administration, Funding acquisition.

## Acknowledgments

We would like to thank dr. C. Moers and his research group for providing the human adipose tissue samples used in this project. Also, the Endothelial Cell Facility of the University of Medical Center Groningen for proving us with the HUVEC used in the TFA experiments. Additionally, we thank dr. M. M. Faas for her all the help and input.

