## Supplementary figure and tables for "Modulating adipose-derived stromal cells’ secretomes by culture conditions: effects on angiogenesis, collagen deposition, and immunomodulation"

**SUPPLEMENTARY MATERIAL**

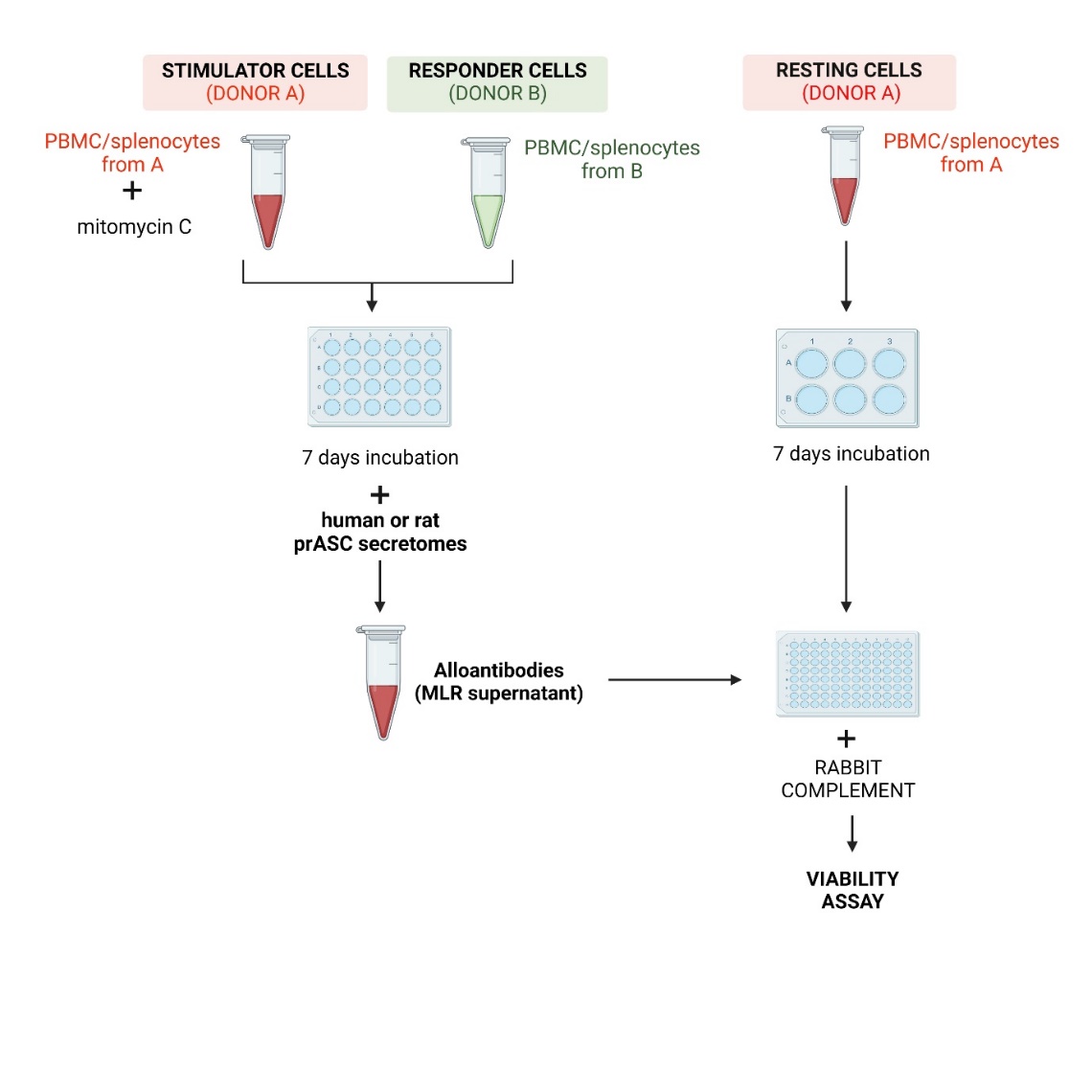
**Supplementary Figure 1.** Illustrative representation of the one-way mixed lymphocyte reaction (MLR) protocol used to evaluate the various human and rat prASC secretomes' capacity to modulate antibody-mediate immune responses.

**DESCRIPTIVE STATISTICS**

**Supplementary table 1.** Detailed overview of the statistical results obtained in the tube formation assay using h-prASC secretomes.

| Dunn's multiple comparisons test | Mean rank diff, | 95,00% CI of diff, | Summary | Adjusted P Value |
| --- | --- | --- | --- | --- |
| STD-M (-) vs. Normoxia | -1,239 | -1,439 to -1,038 | **** | <0,0001 |
| STD-M (-) vs. Cytokines | -0,5872 | -0,7703 to -0,4041 | *** | 0,0006 |
| STD-M (-) vs. High Glucose | -0,7788 | -1,172 to -0,3858 | ** | 0,0043 |
| STD-M (-) vs. Hypoxia | -1,439 | -1,810 to -1,069 | *** | 0,0003 |
| STD-M (-) vs. Hypoxia + High Glucose | -0,7372 | -1,167 to -0,3075 | ** | 0,0073 |
| Normoxia vs. Cytokines | 0,6516 | 0,4161 to 0,8871 | ** | 0,0012 |
| Normoxia vs. High Glucose | 0,4600 | 0,05148 to 0,8685 | * | 0,0335 |
| Normoxia vs. Hypoxia | -0,2006 | -0,6480 to 0,2468 | ns | 0,4178 |
| Normoxia vs. Hypoxia + High Glucose | 0,5016 | 0,05654 to 0,9467 | * | 0,0334 |
| Cytokines vs. High Glucose | -0,1916 | -0,4306 to 0,04735 | ns | 0,1006 |
| Cytokines vs. Hypoxia | -0,8522 | -1,133 to -0,5709 | *** | 0,0008 |
| Cytokines vs. Hypoxia + High Glucose | -0,1500 | -0,6947 to 0,3947 | ns | 0,7736 |
| High Glucose vs. Hypoxia | -0,6606 | -0,8805 to -0,4407 | *** | 0,0008 |
| High Glucose vs. Hypoxia + High Glucose | 0,04160 | -0,6653 to 0,7485 | ns | 0,9996 |
| Hypoxia vs. Hypoxia + High Glucose | 0,7022 | 0,1114 to 1,293 | * | 0,0278 |

|  | STD-M (-) | Normoxia | Cytokines | High Glucose | Hypoxia | Hypoxia + High Glucose |
| --- | --- | --- | --- | --- | --- | --- |
| Number of values | 5 | 5 | 5 | 5 | 5 | 5 |
| Missing values | 0 | 0 | 0 | 0 | 0 | 0 |
| Minimum | 1,000 | 2,120 | 1,489 | 1,500 | 2,138 | 1,471 |
| 25% Percentile | 1,000 | 2,150 | 1,519 | 1,601 | 2,289 | 1,563 |
| Median | 1,000 | 2,228 | 1,556 | 1,813 | 2,516 | 1,735 |
| 75% Percentile | 1,000 | 2,334 | 1,672 | 1,940 | 2,552 | 1,913 |
| Maximum | 1,000 | 2,339 | 1,713 | 1,961 | 2,567 | 2,024 |
| Mean | 1,000 | 2,239 | 1,587 | 1,779 | 2,439 | 1,737 |
| Std. Deviation | 0,000 | 0,09461 | 0,08633 | 0,1853 | 0,1749 | 0,2026 |
| Std. Error of Mean | 0,000 | 0,04231 | 0,03861 | 0,08288 | 0,07821 | 0,09062 |
| Lower 95% CI | 1,000 | 2,121 | 1,480 | 1,549 | 2,222 | 1,486 |
| Upper 95% CI | 1,000 | 2,356 | 1,694 | 2,009 | 2,657 | 1,989 |

**Supplementary table 2.** Detailed overview of the statistical results obtained in the tube formation assay using r-prASC secretomes.

| Tukey's multiple comparisons test | Mean Diff, | 95,00% CI of diff, | Summary | Adjusted P Value |
| --- | --- | --- | --- | --- |
| STD-M (-) vs. Normoxia | -0,9400 | -1,296 to -0,5843 | ** | 0,0031 |
| STD-M (-) vs. Cytokines | -0,2850 | -0,4173 to -0,1527 | ** | 0,0056 |
| STD-M (-) vs. High Glucose | -0,3925 | -0,8052 to 0,02018 | ns | 0,0572 |
| STD-M (-) vs. Hypoxia | -1,395 | -1,948 to -0,8418 | ** | 0,0035 |
| STD-M (-) vs. Hypoxia + High Glucose | -0,6450 | -1,032 to -0,2578 | * | 0,0120 |
| Normoxia vs. Cytokines | 0,6550 | 0,2678 to 1,042 | * | 0,0114 |
| Normoxia vs. High Glucose | 0,5475 | 0,1576 to 0,9374 | * | 0,0195 |
| Normoxia vs. Hypoxia | -0,4550 | -1,363 to 0,4533 | ns | 0,2647 |
| Normoxia vs. Hypoxia + High Glucose | 0,2950 | -0,2815 to 0,8715 | ns | 0,2535 |
| Cytokines vs. High Glucose | -0,1075 | -0,4812 to 0,2662 | ns | 0,6324 |
| Cytokines vs. Hypoxia | -1,110 | -1,680 to -0,5398 | ** | 0,0076 |
| Cytokines vs. Hypoxia + High Glucose | -0,3600 | -0,6413 to -0,07870 | * | 0,0253 |
| High Glucose vs. Hypoxia | -1,003 | -1,848 to -0,1572 | * | 0,0313 |
| High Glucose vs. Hypoxia + High Glucose | -0,2525 | -0,5850 to 0,07999 | ns | 0,1024 |
| Hypoxia vs. Hypoxia + High Glucose | 0,7500 | 0,1219 to 1,378 | * | 0,0308 |

|  | STD-M (-) | Normoxia | Cytokines | High Glucose | Hypoxia | Hypoxia + High Glucose |
| --- | --- | --- | --- | --- | --- | --- |
| Number of values | 4 | 4 | 4 | 4 | 4 | 4 |
| Missing values | 0 | 0 | 0 | 0 | 0 | 0 |
| Minimum | 1,000 | 1,760 | 1,240 | 1,250 | 2,270 | 1,530 |
| 25% Percentile | 1,000 | 1,808 | 1,248 | 1,260 | 2,270 | 1,543 |
| Median | 1,000 | 1,985 | 1,275 | 1,385 | 2,315 | 1,605 |
| 75% Percentile | 1,000 | 2,028 | 1,333 | 1,533 | 2,600 | 1,788 |
| Maximum | 1,000 | 2,030 | 1,350 | 1,550 | 2,680 | 1,840 |
| Mean | 1,000 | 1,940 | 1,285 | 1,393 | 2,395 | 1,645 |
| Std. Deviation | 0,000 | 0,1252 | 0,04655 | 0,1452 | 0,1947 | 0,1363 |
| Std. Error of Mean | 0,000 | 0,06258 | 0,02327 | 0,07261 | 0,09734 | 0,06813 |
| Lower 95% CI | 1,000 | 1,741 | 1,211 | 1,161 | 2,085 | 1,428 |
| Upper 95% CI | 1,000 | 2,139 | 1,359 | 1,624 | 2,705 | 1,862 |

**Supplementary table 3.** Detailed overview of the statistical results obtained in the Sirius-red staining assay using h-prASC secretomes.

| Tukey's multiple comparisons test | Mean Diff, | 95,00% CI of diff, | Summary | Adjusted P Value |
| --- | --- | --- | --- | --- |
| STD-M (-) vs. Normoxia | -0,4723 | -0,6931 to -0,2514 | ** | 0,0058 |
| STD-M (-) vs. Cytokines | 0,1065 | -0,03328 to 0,2463 | ns | 0,1016 |
| STD-M (-) vs. High Glucose | 0,3043 | 0,1267 to 0,4818 | * | 0,0110 |
| STD-M (-) vs. Hypoxia | -0,6425 | -0,8341 to -0,4509 | ** | 0,0016 |
| STD-M (-) vs. Hypoxia + High Glucose | -0,3000 | -0,5838 to -0,01618 | * | 0,0430 |
| Normoxia vs. Cytokines | 0,5788 | 0,4411 to 0,7164 | *** | 0,0010 |
| Normoxia vs. High Glucose | 0,7765 | 0,5173 to 1,036 | ** | 0,0022 |
| Normoxia vs. Hypoxia | -0,1703 | -0,4342 to 0,09374 | ns | 0,1522 |
| Normoxia vs. Hypoxia + High Glucose | 0,1723 | 0,04947 to 0,2950 | * | 0,0195 |
| Cytokines vs. High Glucose | 0,1978 | 0,06658 to 0,3289 | * | 0,0159 |
| Cytokines vs. Hypoxia | -0,7490 | -0,8848 to -0,6132 | *** | 0,0004 |
| Cytokines vs. Hypoxia + High Glucose | -0,4065 | -0,5769 to -0,2361 | ** | 0,0041 |
| High Glucose vs. Hypoxia | -0,9468 | -0,9635 to -0,9300 | **** | <0,0001 |
| High Glucose vs. Hypoxia + High Glucose | -0,6043 | -0,8488 to -0,3597 | ** | 0,0037 |
| Hypoxia vs. Hypoxia + High Glucose | 0,3425 | 0,09764 to 0,5874 | * | 0,0197 |

|  | STD-M (-) | Normoxia | Cytokines | High Glucose | Hypoxia | Hypoxia + High Glucose |
| --- | --- | --- | --- | --- | --- | --- |
| Number of values | 4 | 4 | 4 | 4 | 4 | 4 |
| Missing values | 0 | 0 | 0 | 0 | 0 | 0 |
| Minimum | 1,000 | 1,373 | 0,8480 | 0,6060 | 1,547 | 1,213 |
| 25% Percentile | 1,000 | 1,392 | 0,8505 | 0,6305 | 1,573 | 1,214 |
| Median | 1,000 | 1,486 | 0,8870 | 0,7145 | 1,660 | 1,292 |
| 75% Percentile | 1,000 | 1,539 | 0,9430 | 0,7423 | 1,695 | 1,395 |
| Maximum | 1,000 | 1,544 | 0,9520 | 0,7480 | 1,703 | 1,404 |
| Mean | 1,000 | 1,472 | 0,8935 | 0,6958 | 1,643 | 1,300 |
| Std. Deviation | 0,000 | 0,07771 | 0,04919 | 0,06247 | 0,06742 | 0,09988 |
| Std. Error of Mean | 0,000 | 0,03885 | 0,02460 | 0,03124 | 0,03371 | 0,04994 |
| Lower 95% CI | 1,000 | 1,349 | 0,8152 | 0,5963 | 1,535 | 1,141 |
| Upper 95% CI | 1,000 | 1,596 | 0,9718 | 0,7952 | 1,750 | 1,459 |

**Supplementary table 4.** Detailed overview of the statistical results obtained in the Sirius-red staining assay using r-prASC secretomes.

| Tukey's multiple comparisons test | Mean Diff, | 95,00% CI of diff, | Summary | Adjusted P Value |
| --- | --- | --- | --- | --- |
| STD-M (-) vs. Normoxia | -0,6808 | -1,256 to -0,1053 | * | 0,0316 |
| STD-M (-) vs. Cytokines | -0,2090 | -0,6296 to 0,2116 | ns | 0,2691 |
| STD-M (-) vs. High Glucose | -0,2663 | -0,7019 to 0,1694 | ns | 0,1724 |
| STD-M (-) vs. Hypoxia | -1,180 | -1,892 to -0,4677 | * | 0,0121 |
| STD-M (-) vs. Hypoxia + High Glucose | -0,3328 | -0,8028 to 0,1373 | ns | 0,1218 |
| Normoxia vs. Cytokines | 0,4718 | 0,3017 to 0,6418 | ** | 0,0027 |
| Normoxia vs. High Glucose | 0,4145 | 0,2356 to 0,5934 | ** | 0,0045 |
| Normoxia vs. Hypoxia | -0,4993 | -0,6820 to -0,3165 | ** | 0,0028 |
| Normoxia vs. Hypoxia + High Glucose | 0,3480 | 0,1547 to 0,5413 | ** | 0,0095 |
| Cytokines vs. High Glucose | -0,05725 | -0,2053 to 0,09084 | ns | 0,4245 |
| Cytokines vs. Hypoxia | -0,9710 | -1,294 to -0,6478 | ** | 0,0021 |
| Cytokines vs. Hypoxia + High Glucose | -0,1238 | -0,2668 to 0,01926 | ns | 0,0734 |
| High Glucose vs. Hypoxia | -0,9138 | -1,202 to -0,6251 | ** | 0,0019 |
| High Glucose vs. Hypoxia + High Glucose | -0,06650 | -0,2083 to 0,07528 | ns | 0,3015 |
| Hypoxia vs. Hypoxia + High Glucose | 0,8473 | 0,5869 to 1,108 | ** | 0,0018 |

|  | STD-M (-) | Normoxia | Cytokines | High Glucose | Hypoxia | Hypoxia + High Glucose |
| --- | --- | --- | --- | --- | --- | --- |
| Number of values | 4 | 4 | 4 | 4 | 4 | 4 |
| Missing values | 0 | 0 | 0 | 0 | 0 | 0 |
| Minimum | 1,000 | 1,498 | 1,080 | 1,102 | 1,953 | 1,169 |
| 25% Percentile | 1,000 | 1,507 | 1,086 | 1,120 | 1,960 | 1,180 |
| Median | 1,000 | 1,648 | 1,180 | 1,270 | 2,162 | 1,334 |
| 75% Percentile | 1,000 | 1,888 | 1,362 | 1,410 | 2,419 | 1,484 |
| Maximum | 1,000 | 1,930 | 1,397 | 1,424 | 2,444 | 1,494 |
| Mean | 1,000 | 1,681 | 1,209 | 1,266 | 2,180 | 1,333 |
| Std. Deviation | 0,000 | 0,2025 | 0,1480 | 0,1533 | 0,2507 | 0,1654 |
| Std. Error of Mean | 0,000 | 0,1013 | 0,07401 | 0,07665 | 0,1253 | 0,08271 |
| Lower 95% CI | 1,000 | 1,358 | 0,9735 | 1,022 | 1,781 | 1,070 |
| Upper 95% CI | 1,000 | 2,003 | 1,445 | 1,510 | 2,579 | 1,596 |

**Supplementary table 5.** Detailed overview of the statistical results obtained in the mixed lymphocyte reaction (MLR) assay using h-prASC secretomes.

| Tukey's multiple comparisons test | Mean Diff. | 95,00% CI of diff. | Summary | Adjusted P Value |
| --- | --- | --- | --- | --- |
| MLR supernatant vs. Normoxia | -231,4 | -273,4 to -189,5 | **** | <0,0001 |
| MLR supernatant vs. Cytokines | -62,38 | -103,1 to -21,61 | ** | 0,0039 |
| MLR supernatant vs. High Glucose | -4,420 | -48,51 to 39,67 | ns | 0,9990 |
| MLR supernatant vs. Hypoxia | -214,9 | -251,7 to -178,1 | **** | <0,0001 |
| MLR supernatant vs. Hypoxia + High Glucose | -139,6 | -172,6 to -106,5 | **** | <0,0001 |
| Normoxia vs. Cytokines | 169,1 | 97,22 to 240,9 | *** | 0,0002 |
| Normoxia vs. High Glucose | 227,0 | 158,6 to 295,5 | **** | <0,0001 |
| Normoxia vs. Hypoxia | 16,51 | -52,69 to 85,72 | ns | 0,9502 |
| Normoxia vs. Hypoxia + High Glucose | 91,86 | 45,52 to 138,2 | *** | 0,0006 |
| Cytokines vs. High Glucose | 57,96 | -4,310 to 120,2 | ns | 0,0711 |
| Cytokines vs. Hypoxia | -152,5 | -210,7 to -94,40 | **** | <0,0001 |
| Cytokines vs. Hypoxia + High Glucose | -77,20 | -133,1 to -21,29 | ** | 0,0077 |
| High Glucose vs. Hypoxia | -210,5 | -267,1 to -153,9 | **** | <0,0001 |
| High Glucose vs. Hypoxia + High Glucose | -135,2 | -205,6 to -64,70 | *** | 0,0008 |
| Hypoxia vs. Hypoxia + High Glucose | 75,35 | 36,40 to 114,3 | *** | 0,0007 |

|  | STD-M (-) | Normoxia | Cytokines | High Glucose | Hypoxia | Hypoxia + High Glucose |
| --- | --- | --- | --- | --- | --- | --- |
| Number of values | 10 | 10 | 10 | 10 | 10 | 10 |
| Missing values | 0 | 0 | 0 | 0 | 0 | 0 |
| Minimum | 100,0 | 258,8 | 94,70 | 54,48 | 271,6 | 201,6 |
| 25% Percentile | 100,0 | 311,7 | 132,9 | 74,25 | 285,3 | 212,5 |
| Median | 100,0 | 329,4 | 174,4 | 92,60 | 316,1 | 233,6 |
| 75% Percentile | 100,0 | 358,5 | 188,7 | 145,9 | 340,5 | 268,9 |
| Maximum | 100,0 | 392,3 | 200,4 | 172,4 | 369,1 | 283,0 |
| Mean | 100,0 | 331,4 | 162,4 | 104,4 | 314,9 | 239,6 |
| Std. Deviation | 0,000 | 37,36 | 36,29 | 39,25 | 32,75 | 29,41 |
| Std. Error of Mean | 0,000 | 11,81 | 11,48 | 12,41 | 10,36 | 9,300 |
| Lower 95% CI | 100,0 | 304,7 | 136,4 | 76,34 | 291,5 | 218,5 |
| Upper 95% CI | 100,0 | 358,2 | 188,3 | 132,5 | 338,3 | 260,6 |

**Supplementary table 6.** Detailed overview of the statistical results obtained in the mixed lymphocyte reaction (MLR) assay using r-prASC secretomes.

| Tukey's multiple comparisons test | Mean Diff, | 95,00% CI of diff, | Summary | Adjusted P Value |
| --- | --- | --- | --- | --- |
| MLR supernatant vs. Normoxia | -187,8 | -229,1 to -146,6 | **** | <0,0001 |
| MLR supernatant vs. Cytokines | -69,48 | -120,1 to -18,87 | * | 0,0117 |
| MLR supernatant vs. High Glucose | -44,81 | -87,03 to -2,597 | * | 0,0387 |
| MLR supernatant vs. Hypoxia | -142,8 | -189,8 to -95,77 | *** | 0,0002 |
| MLR supernatant vs. Hypoxia + High Glucose | -61,54 | -106,7 to -16,34 | * | 0,0122 |
| Normoxia vs. Cytokines | 118,4 | 33,81 to 202,9 | * | 0,0106 |
| Normoxia vs. High Glucose | 143,0 | 71,32 to 214,7 | ** | 0,0017 |
| Normoxia vs. Hypoxia | 45,05 | 9,621 to 80,47 | * | 0,0169 |
| Normoxia vs. Hypoxia + High Glucose | 126,3 | 64,84 to 187,7 | ** | 0,0014 |
| Cytokines vs. High Glucose | 24,66 | -7,506 to 56,83 | ns | 0,1384 |
| Cytokines vs. Hypoxia | -73,31 | -157,0 to 10,34 | ns | 0,0851 |
| Cytokines vs. Hypoxia + High Glucose | 7,934 | -49,33 to 65,20 | ns | 0,9911 |
| High Glucose vs. Hypoxia | -97,97 | -175,6 to -20,35 | * | 0,0175 |
| High Glucose vs. Hypoxia + High Glucose | -16,73 | -64,55 to 31,09 | ns | 0,7320 |
| Hypoxia vs. Hypoxia + High Glucose | 81,24 | 24,60 to 137,9 | ** | 0,0094 |

|  | STD-M (-) | Normoxia | Cytokines | High Glucose | Hypoxia | Hypoxia + High Glucose |
| --- | --- | --- | --- | --- | --- | --- |
| Number of values | 7 | 7 | 7 | 7 | 7 | 7 |
| Missing values | 0 | 0 | 0 | 0 | 0 | 0 |
| Minimum | 100,0 | 232,4 | 116,2 | 112,8 | 198,3 | 108,4 |
| 25% Percentile | 100,0 | 278,4 | 150,3 | 122,3 | 211,1 | 142,8 |
| Median | 100,0 | 292,3 | 166,7 | 129,7 | 246,7 | 157,5 |
| 75% Percentile | 100,0 | 309,8 | 188,7 | 176,3 | 270,9 | 188,9 |
| Maximum | 100,0 | 312,8 | 225,0 | 182,1 | 285,6 | 192,6 |
| Mean | 100,0 | 287,8 | 169,5 | 144,8 | 242,8 | 161,5 |
| Std. Deviation | 0,000 | 27,43 | 33,64 | 28,06 | 31,25 | 30,05 |
| Std. Error of Mean | 0,000 | 10,37 | 12,72 | 10,61 | 11,81 | 11,36 |
| Lower 95% CI | 100,0 | 262,5 | 138,4 | 118,9 | 213,9 | 133,8 |
| Upper 95% CI | 100,0 | 313,2 | 200,6 | 170,8 | 271,7 | 189,3 |
